# Imaging of red-shifted photons from bioluminescent tumours using fluorescence by unbound excitation from luminescence

**DOI:** 10.1101/428771

**Authors:** Fabiane Sônego, Sophie Bouccara, Thomas Pons, Nicolas Lequeux, Anne Danckaert, Jean-Yves Tinevez, Israt S. Alam, Spencer L. Shorte, Régis Tournebize

## Abstract

Early detection of tumours is today a major challenge and requires sensitive imaging methodologies coupled with new efficient probes. Bioluminescence imaging has been widely used in the field of oncology and several cancer cell lines have been genetically modified to provide bioluminescence signals. However, photons that are emitted by the majority of commonly used luciferases are usually in the blue part of the visible spectrum, where tissue absorption is still very high, making deep tissue imaging non-optimal and calling for optimised optical imaging methodologies. We have previously shown that red-shifting of bioluminescence signal by Fluorescence Unbound Excitation from Luminescence (FUEL) is a mean to increase bioluminescence signal sensitivity detection *in vivo*. Here, we applied FUEL to tumour detection in two different subcutaneous tumour models: the auto-luminescent human embryonic kidney (HEK293) cell line and the murine B16-F10 melanoma cell line previously transfected with the plasmid Luc2. Tumour size and bioluminescence were measured over time and tumour vascularization characterized. We then locally injected near infrared emitting Quantum Dots (NIR QDs)in the tumour site and observed a red-shifting of bioluminescence signal by (FUEL) indicating that FUEL could be used to allow deeper tumour detection.

## Introduction

Imaging of physiological and pathological processes benefits from sensitive methodologies [1] and new imaging probes and methodologies are constantly evolving from the progress in preclinical research and important insights that it has yielded. Preclinical and small-animal imaging modalities allow longitudinal and multiparametric studies while reducing the number of animals used in the studies and thus comply with ethical guidelines. They include MRI, SPECT, and PET [1, 2]. Whilst MRI and nuclear imaging confer high resolution and sensitivity respectively, the cost of these scanners and their maintenance represent major limitations in their use. By contrast, optical imaging is a widely used and low-cost methodology, also offering high sensitivity but also high throughput [3].

Bioluminescence imaging has been widely used in the field of oncology. Several cell lines have been genetically modified to provide both *in vitro* and *in vivo* stable bioluminescence signals. In most cases, tumour cells are modified to express the enzyme luciferase and then a suitable substrate is added exogenously, which leads to the production of light in presence of oxygen and ATP [3, 4]. Recently, autonomous bioluminescent mammalian cell lines have been developed. These cell lines express both codon-optimised *Photorhabdus luminescens* luciferases coding genes and associated genes responsible for the production and recycling of aldehyde and FMNH_2_ co-substrates required for light emission. As a direct consequence, these cell lines do not require substrate addition to be luminescent [5]. Photon production in bioluminescence is chemically dependent, provides high sensitivity and low background signals, and unlike fluorescence does not require external excitation sources. However, the optical spectral region where luciferases maximally emit is between 480 and 620 nm, where tissue absorption is maximum, highly limiting deep tissue bioluminescence imaging [6, 3] while a range of wavelengths between 650 and 900 nm is more suitable for *in vivo* imaging [7]. Several strategies have been developed in the last few years to overcome this limitation by red-shifting the emission in the well-adapted wavelength range where tissue absorption is minimal. One of the strategies adopted is the Bioluminescence Resonance Energy Transfer (BRET). BRET is a non-radiative process in which energy is transferred from a bioluminescent donor to a fluorescent acceptor that has been shown to be a powerful tool to evaluate protein-protein interaction [8, 9]. Based on the principle of BRET, self-illuminated quantum dots (QDs) have been designed [10]. QDs are inorganic fluorescent nanocrystals that are ideal candidate as BRET acceptor due to their broad absorbance spectra, high absorbance cross sections, high fluorescence quantum yield and their large Stokes shift in the near infrared (NIR) region [11]. In this context, carboxylate QDs coupled with amide luciferase and even functionalized with a RGD peptide have been developed for targeting *in vivo* cancer cells [12–14].

Recently, we reported Fluorescence by Unbound Excitation from Luminescence (FUEL) as a mean to red-shift bioluminescence emission without requiring extremely close contact between donor and acceptor like in BRET. FUEL is defined as a radiative transfer between a bioluminescent source exciting nearby fluorophore [15, 16]. We have hypothesized that FUEL could be a useful tool for the detection of tumours *in vivo* due to two main advantages. Firstly, luciferase does not need to be grafted to the nanoparticles. This would allow the use of smaller diameter nanoparticles, likely to have superior pharmacokinetic properties in comparison to coupled larger nanoparticles [17, 18]. Secondly, because in FUEL, QDs red-emission is spatially correlated with the bioluminescence emission of tumour cells, it is a relevant mean to increase the sensitivity of the signal in tissue and is a marker of proximity.

In this study, we used two different *in vivo* subcutaneous bioluminescent tumour models to investigate the suitability of FUEL in detecting tumours. The first model was induced by bioluminescent B16-F10 tumour cells expressing firefly luciferase [19–21]. These cells will be referred here as B16-Luc2. The second tumour model established here was a bioluminescent HEK293 model, a human embryonic kidney cell line expressing the lux operon from bacteria and will hereon be referred as HEK-Lux. This cell type expresses both the luciferase and enzymes required for the production of the substrate, and therefore does not require further administration of substrate [5]. Using these two models, we present and quantify the first *in vivo* FUEL experiments using near-infrared emitting quantum dots to achieve a red-shifting emission of the subcutaneous tumours.

## Methods

### Cell lines culture

The autobioluminescent HEK293 cells with the luxCDABE operon (HEK-Lux) cells were kindly provided by 490 BioTech (Tennessee, USA)[22]. These cells were cultured at 37°C and 5% CO_2_ in DMEM with Glutamax and Pyruvate (Life technologies) supplemented with 10% heat-inactivated fetal bovine serum (FBS, Gibco), 1% of non-essential amino acids (Sigma), 1% penicillin/streptomycin (Life technologies) and 100 μg/mL G418 (Sigma). The experiments were performed with cells at passage 20 to 22. Non-autobioluminescent HEK293 cells were cultured in the same medium as HEK-Lux cells, but in the absence of antibiotic G418. At confluence, cells were rinsed with phosphate buffered saline without Ca^2+^ and Mg^2+^ (PBS, Gibco) and harvested with 0.05% trypsin-EDTA (Gibco). Cells were used at passage 9.

The melanoma cell line B16-F10, expressing Luc2 (B16-Luc2) was kindly provided by the group of Pierre Bruhns (Institut Pasteur, Paris). The cells were cultured in RPMI 1640 with glutamine and Hepes (Gibco) supplemented with 10% heat-inactivated FBS and 1% penicillin/streptomycin. At maximum 50% of confluence, cells were rinsed with PBS and harvested with 0.05% trypsin-EDTA. The experiments were performed with cells at passages between 6 and 16.

The emission spectra of the HEK-Lux and B16-Luc2 cells were determined using 2×10^5^ cells suspended in 0.1 mL of appropriated medium. One day prior to imaging, cells were seeded in a 96-well clear bottom black plate (Nunc) and incubated overnight at 37°C and 5% CO2. The medium was gently removed from the wells and replaced with fresh medium prior to image acquisition. For B16-Luc2 cells, the substrate D-luciferin (Perkin Elmer) was added to the cells (150 μg/mL in 0.01 mL). Bioluminescence images were acquired with an IVIS Spectrum system, using 20 nm bandpass emission filters and OPEN mode (exposure time of 180 sec for HEK-Lux cells and 30 sec for B16-Luc2 cells).

### Mice and ethics statement

Female nude mice (Rj:NMRI-nu) (7 weeks-old) were obtained from Janvier Laboratories (France). All protocols involving animal experiments were approved and carried out in accordance with the ethical guidelines of Institut Pasteur, Paris (license number: 2014-0055). The mice were housed in the Biosafety Level 2+ animal facility of Institut Pasteur. All mice had free access to food and water and were under controlled light/dark cycle, temperature and humidity. Animals were handled with regard for alleviation of suffering. Animals were anesthetized using isoflurane, and euthanized with CO2.

### Induction of subcutaneous tumours

#### HEK-Lux and non-bioluminescent HEK models

Each tumour was induced by subcutaneous (s.c.) administration of 0.1 mL of 5×10^6^ cells (suspended in medium without FBS) and basement membrane matrix growth factor reduced (matrigel Corning), (25:75, v/v).

#### B16-Luc2 model

Each tumour was induced by s.c. administration of 0.1 mL of 8×10^4^ cells (suspended in medium without FBS) and basement membrane matrix growth factor reduced (matrigel, Corning), (20:80, v/v).

For all cell lines, culture medium was replaced with fresh medium one day prior to the subcutaneous injection.

Two ventral tumours were induced in each mouse. The mice were anesthetized with 2% isoflurane gas prior to the injection of the tumour cells. Cells were first administered subcutaneously on the left side and then on the right side of the mice. All the results shown here represent measurements taken for the left tumour of each mouse. Tumour growth was monitored by calliper measurement and determined as previously described; volume = [(width/2)^2^ × length] [23].

### Near infra-red (NIR) QDs

NIR QDs were synthesized as previously described [24] and water-solubilized as described in [25]. NIR QDs were diluted in PBS to provide the desired concentration. Absorption and emission spectra of a 0.1 μM solution were determined using IVIS Spectrum.

### *In vivo* bioluminescence and fluorescence imaging

Bioluminescence and fluorescence imaging were performed using an IVIS Spectrum system (Perkin Elmer). Unless specified elsewhere, mice bearing the autobioluminescent HEK-Lux tumours were anesthetized with 2% isoflurane gas and typically imaged with (840 nm) and without emission filter (total light output - open filter) for 300 sec. Mice bearing the bioluminescent B16-Luc2 tumours were intraperitoneally (i.p.) administered with the substrate D-luciferin (0.75 mg/mouse, Perkin Elmer) 11 min prior to bioluminescence imaging. This time point was chosen to allow a comparison between different mice and because it corresponds to the D-luciferin peak bioavailability. Mice were anesthetized with 2% isoflurane gas immediately after the administration of D-luciferin and maintained under anesthesia until the end of the image acquisition. Bioluminescence images were acquired in the open mode or with the 840 nm filter for 180, 60 or 3 sec, as specified in figures legends. Fluorescence images were also acquired using IVIS Spectrum system (excitation filter 430 nm and emission filter 840 nm +/− 20 nm). Living Image software (Perkin Elmer) was used to define and analyse the light emission in the regions of interest (ROIs). *Angiosense 750EX:* The fluorescent vascular agent Angiosense 750EX (Perkin Elmer) was administered intravenously (i.v.)(2 nmol/0.1 mL) in mice bearing HEK-Lux or B16-Luc2 tumours, 22 to 30 or 7 to 9 days post tumour cells injection, respectively. Mice were anesthetized with 2% isoflurane gas prior to the image acquisition. The vascularization of the tumours was evaluated 24 h post Angiosense 750EX administration using the IVIS Spectrum system. Fluorescent images were acquired with 745 nm excitation filter and 800 nm emission filter, with the auto option selected as time of exposure.

#### NIR QDs

Fluorescent images using IVIS Spectrum were acquired prior and after NIR QDs intratumoral administration *in vivo* with 0.1 sec of exposure time, and 430 and 840 nm as excitation and emission filters, respectively.

### Dextran- Fluorescein isothiocyanate (FITC)

High molecular weight dextran-FITC (500 KMW, Molecular Probes) was injected i.v. *via* the retro-orbital sinus (0.5 mg/0.1 mL) in mice bearing HEK-Lux or B16-Luc2 tumours. Harvested tumours were fixed in 4% parafolmadehyde (EMC) for 3 to 5 hours at room temperature, depending on the tumour volume, followed by aldehydes quenching by 1 h incubation in 100mM glycine (Sigma-Aldrich). Tumours were then incubated in 15% sucrose (Sigma-Aldrich) at 4°C overnight and in 30% sucrose at 4°C for approximately 24 h before embedding in Shandon Cryomatrix (Thermo Fischer) and freezing using isopentanol. Fifty μm sections cut using cryostat (CM3050 S, Leica) were stained with DAPI and imaged using an automated spinning disk microscope CellVoyager1000 (Yokogawa Electrics, Japan). The sections were left overnight at room temperature before being stained with DAPI.

### FUEL experiments

#### In vitro FUEL

B16-Luc2, HEK-Lux and HEK non-bioluminescent cells (2×10^5^, 0.1 mL of appropriated medium) were seeded in a 96-well clear bottom black plate (Nunc) one day prior to the experiment and incubated at 37°C and 5% CO_2_. On the day of the experiment, the medium was removed and a fresh medium with or without NIR QDs (450 μM in 0.01 mL) was added to the well. Each cell type was cultured with the same medium used for the cell culture. HEK non-bioluminescent cell type was used in this experiment as a negative control for HEK-Lux cells. For B16-Luc2 cells, the substrate D-luciferin was added to the wells (150 μg/mL in 0.01 mL), and the absence of the substrate in the well was used as a negative control for this cell type. Bioluminescence images were acquired with both 840 nm and open filter (exposure time of 300 sec for HEK cells and 180 sec for B16-Luc2 cells). Fluorescence images were also acquired (excitation 430 nm and emission 840 nm, 1 sec as exposure time).

#### Experiments with mice bearing B16-Luc2 tumours

In order to evaluate the bioluminescence signal emitted at 840 nm before the administration of NIR QDs, D-luciferin (0.75 mg/mouse, i.p.) was administered in mice bearing B16-Luc2 tumours 11 min prior to the image acquisition (180 sec as exposure time). After 1 h, bioluminescent images were acquired again to determine the basal bioluminescent signal at 840 nm. Next, 0.5 nmol (0.04 mL) NIR QDs were administered into the left tumour and 0.04 mL PBS into the right tumour. Fluorescence images were acquired (excitation 430 nm/ emission 840 nm, 0.1 sec) prior and post NIR QDs intratumoral administration. D-luciferin was then administered 11 min prior to the bioluminescence imaging acquisition with a 840 nm and open filter for 180 and 3 sec, respectively.

Experiments were also performed to evaluate the possible effect of NIR QDs without a bioluminescence source. For this control, NIR QDs were injected in the left tumour and PBS was injected in the right tumour of the mice, without previous administration of D-luciferin. Both bioluminescence and fluorescence images were acquired, using the same emission and excitation filters and exposure time.

#### Experiments with mice bearing HEK-Lux tumours

Bioluminescence images at 840 nm and open filter (300 sec of exposure time) were acquired prior and post injection of 0.5 nmol (0.04 mL) of NIR QDs in the left tumour and 0.04 mL of PBS in the right tumour of mice bearing the autobioluminescent HEK-Lux tumours. Fluorescence images were acquired (excitation 430 nm and emission 840 nm, 0.1 sec) prior and post NIR QDs intratumoral administration.

### Statistics

The number experimental repeats and animals used for each experiment are noted in the figure legends. When compared, B16-Luc2 and HEK-Lux tumours results were analysed via Mann-Whitney test or Student’s t-test after being assessed for normality of sample distribution. For the statistical analyses, the results from *in vitro* experiments were analysed after normalization by strictly standardized mean difference (SSMD) test as previously described [26]. Statistical analyses and graphs plotting were performed using Prism 6.0 (GraphPad Software Inc.^©^, USA). P-values of *p<0.05 and **p<0.001 were used.

## Results

### Characterisation of tumour models reveals marked differences in bioluminescence emission and growth dynamics but shows similar vascularization

In order to investigate the ability of FUEL to enhance the detection of tumours *in vivo*, we used two distinct bioluminescent preclinical subcutaneous tumour models in nude mice: murine B16-Luc2 melanoma tumours previously described [21] and the human HEK 293 tumor model, adapted from the model described by Ho *et al*. [23].

Firstly, we characterised the emission spectrum for each of the tumoral cell types and observed an emission peak at 600 nm for B16-Luc2 (Fig 1A), while for HEK-Lux the peak was at 500 nm (Fig 1B). It is noteworthy that the B16-Luc2 cells emit a stronger bioluminescent signal when compared to an equal number of HEK-Lux cells. B16-Luc2 cells also showed higher *in vivo* proliferation than HEK-Lux cells. While 8 × 10^4^ B16-Luc2 cells induced the formation of 400 mm^3^ tumours in 14 days (Fig 1C), 5 ×10^6^ HEK-Lux cells were necessary to induced similar tumour sizes in more than 30 days (Fig 1D).

**Fig. 1:**
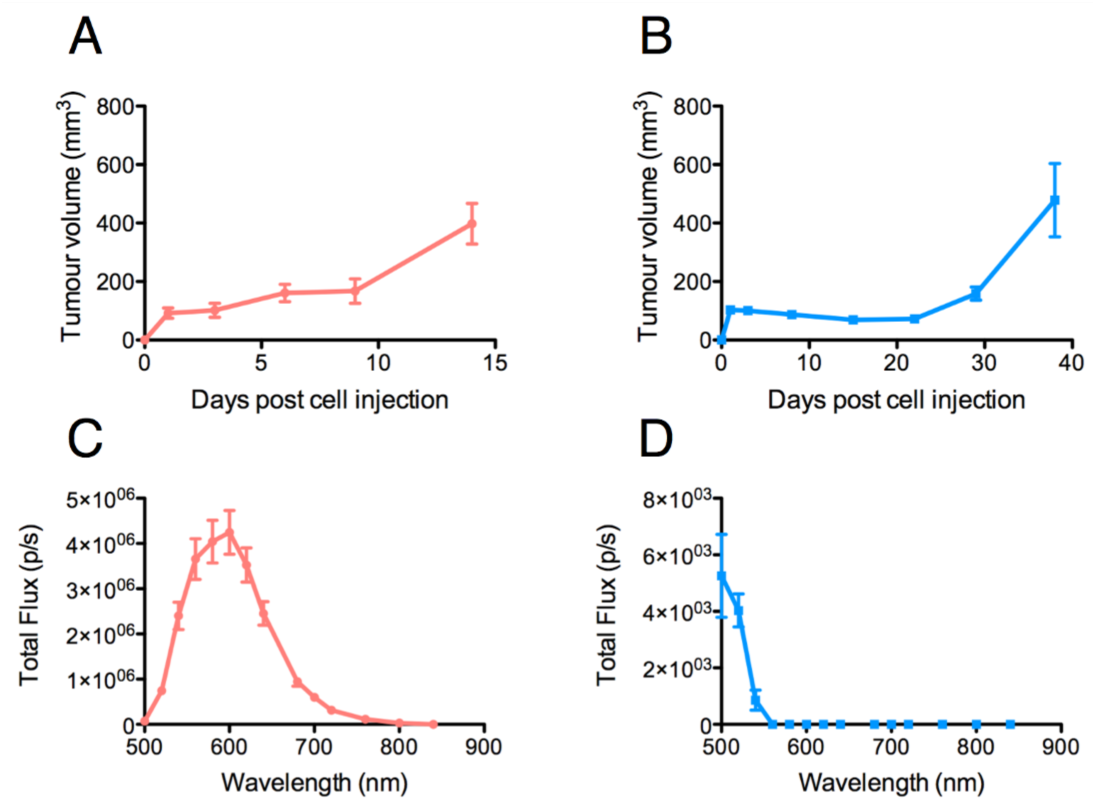
Characterisation of emission spectra of B16-Luc2 and HEK-Lux cells and tumour growth curves. A) Emission spectrum of B16-Luc2 and B) HEK-Lux cells. Bioluminescence images were acquired from 500 to 840 nm for 30 sec (B16-Luc2) or 180 sec (HEK-Lux). Results are expressed as total flux (photons/sec) in the ROI, n=3. C) Tumour growth of B16-Luc2 (8×10^4^, 0.1mL) and D) HEK-Lux (5×10^6^) cells over time, following subcutaneous injection in nude mice on the right and left sides. Results are representative of 4 independent experiments and represent the left tumour volume, n=5. Data shown are means ± SEM.

We also acquired bioluminescence images of tumours over time, and observed that similar to the growth in tumour volume, the bioluminescence signal intensity of B16-Luc2 tumours was detectable as early as 3 days post-injection and increased over time to reach approximately 10^8^ photons emitted/sec per tumour on day 14 (Fig 2A and 2C). In contrast, though HEK-Lux cells emitted a high bioluminescence signal immediately after the subcutaneous injection, this signal disappeared on day 1. The signal stayed low until day 29, when it started to increase again, reaching a maximum of 10^5^ photons/sec per tumour on day 38 (Fig 2B and 2D). Interestingly, the signal increase correlated with the development of the tumour, as assessed by an increase in tumour volume, suggesting that the cells had a latency time before growing and emitting higher bioluminescence signal. Altogether, these observations show that the two tumour models have markedly different growth curves and that the B16-Luc2 tumours emit 1000 times more light using an open filter for detection than the HEK-Lux.

**Fig. 2:**
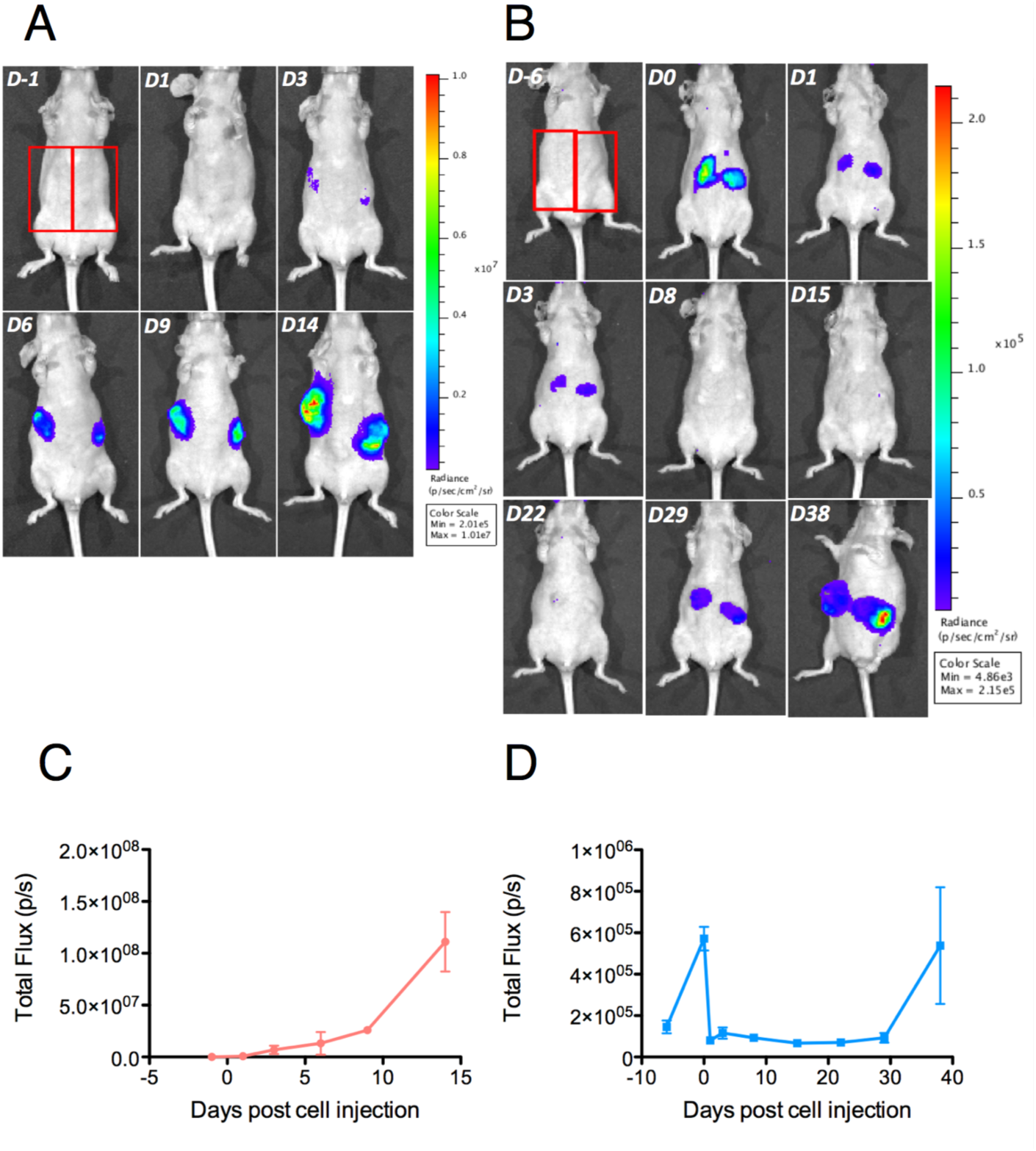
Tumour bioluminescence signal evolution imaging over time. A) B16-Luc2 cells (8×10^4^, 0.1mL) were subcutaneously administered in nude mice. Mice were imaged 1 day prior and 1, 3, 6, 9 and 14 days post administration of B16-Luc2 cells, n=5. B) HEK-Lux cells (5×10^6^, 0.1mL) were subcutaneously adminstered in nude mice. Mice were imaged 6 days prior and 0, 1, 3, 8, 15, 22, 29 and 38 days post administration of HEK-Lux cells, n=6. C) Bioluminescence signal quantitation of B16-Luc2- and D) HEK-Lux-induced tumours. Red rectangles in 2A and 2B show the ROI used for quantification. Results express the total flux (photons/sec) in the ROI of the left tumour of the mice. These results are representative of 4 independent experiments.

We additionally investigated the vascularization of both tumours using the vascular agent Angiosense 750EX. Fluorescence images acquired 24 h post Angiosense administration indicated similar accumulation of the probe in both B16-Luc2 and HEK-Lux-induced tumours (Fig 3A and 3B). Mice not bearing tumours were used as control, and did not show fluorescence signal in the upper abdomen. The fluorescence signal observed in the lower abdomen, in both control and tumour-bearing mice, is likely associated with the renal excretion of the probe. In order to investigate the vascularization at microscopic levels, we have administrated high molecular weight dextran labelled with FITC i.v. Corroborating the results *in vivo*, histological sections suggest that the vascularization is similar in both tumour models (Fig 3C and 3D).

**Fig. 3:**
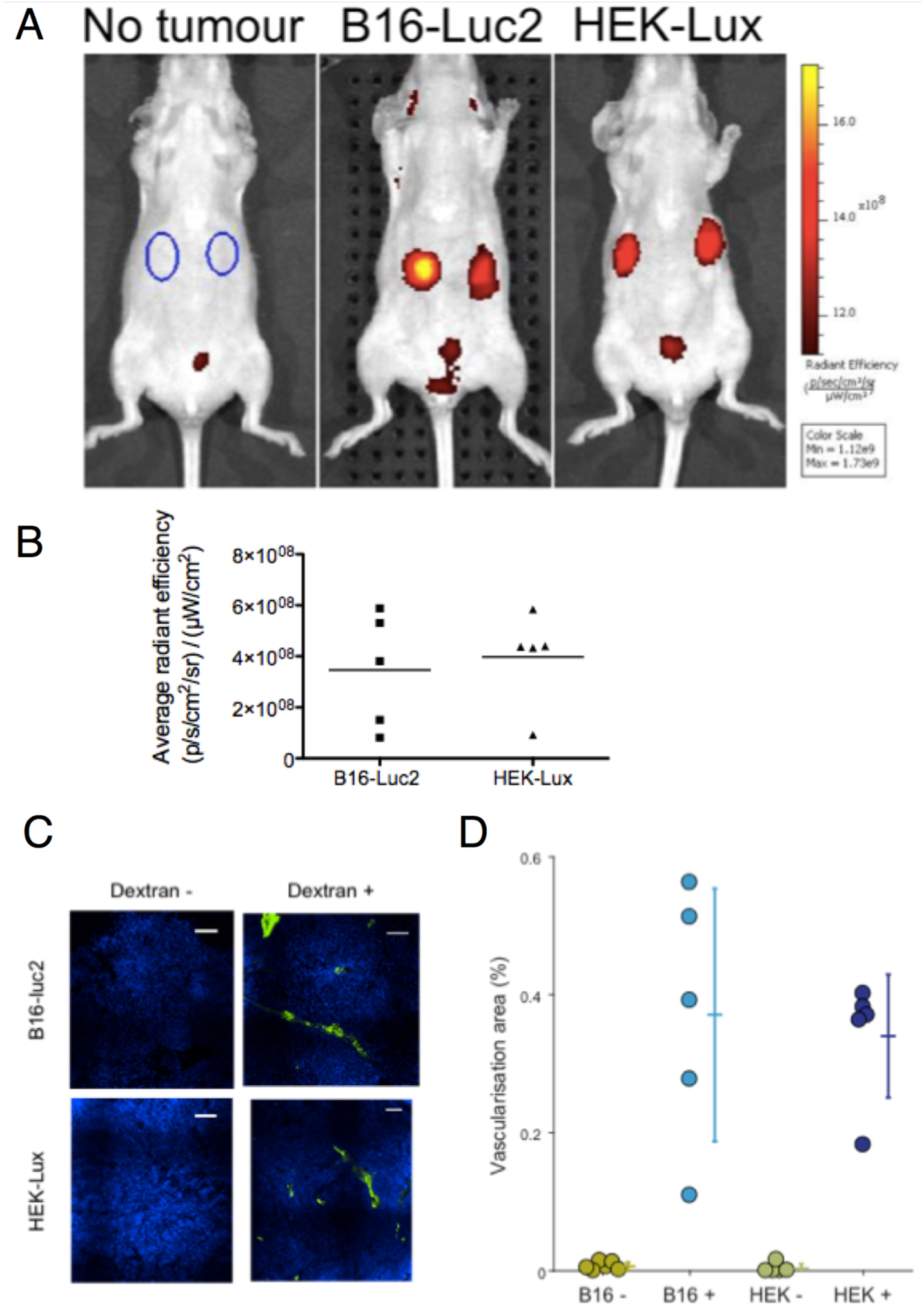
*In vivo* evaluation of tumour vascularisation. A) B16-Luc2 cells (8**×**10^4^, 1mL), HEK-Lux (5 **×** 10^6^, 0.1 mL) were subcutaneously administered in nude mice. Angiosense 750EX (2 nmol, 0.1 mL) was intravenously administered between 7 and 9 days after B16-Luc2 injection or between 22 and 30 days post HEK-Lux cells injection Images were acquired 24 hrs after. B) Fluorescence signal quantitation of Angiosense accumulation in B16-Luc2- and HEK-Lux-induced tumours. ROIs were determined as shown in the first image of Figure 3A. Results express the difference between the average radiant efficiency in the ROI of the left tumour of the mice with tumour and the arithmetic mean of the average radiant efficiency in the ROI of the left side in mice without tumour, (n=4 control group and n=5 for the tumour bearing groups). C) Vizualisation of tumour vascularization using high molecular weight dextran-FITC (500 KMW). Images correspond to a section in the tumors at 50% depth. Contrast and brightness in both channels have been adjusted with an identical color scale across the four images. Scale bars: 100 μm. D) Area of vascularisation, defined as the percentage of the tumour area labelled by dextran at 0, 25, 50, 75 and 100% tumour depth. The area of vascularisation was extracted using an identical threshold over all images.

### FUEL enables enhanced detection of tumours

FUEL efficiency depends on the overlap between the emission spectrum of the bioluminescent source and the excitation spectrum of the acceptor fluorophore. NIR QDs have a broad and continuous decreasing excitation spectrum from UV to 800 nm, as illustrated in Fig 4. This spectrum suggests that both B16-Luc2 (with an emission peak wavelength centred at around 600 nm) and HEK-Lux bioluminescence signal (with an emission peak wavelength centred at around 500 nm) are suitable for the excitation of NIR. Additionally, emission spectrum indicates a maximum emission at around 840 nm. The photoluminescence quantum yield was estimated at 20–30% using ICG in DMSO as a standard fluorophore. Based on these spectra, we first investigated the presence of FUEL with both B16-Luc2 and HEK-Lux *in vitro*. The incubation of B16-Luc2 cells with NIR QDs significantly increased the bioluminescence signal at 840 nm as compared to cells alone, and B16-Luc2 incubated with NIR QDs but in the absence D-luciferin (Fig 4B). Normalized SSMD values classified the FUEL phenomenon extremely strong as compared to the controls (Fig 4C). HEK-Lux cells, which emit weaker bioluminescence signals, also showed an increase in the intensity of bioluminescence at 840 nm in the presence of NIR QDs. The statistical analyses using SSMD normalization indicate a very strong difference between HEK-Lux cells incubated with NIR QDs and controls (HEK-Lux cells alone, and non-bioluminescent HEK cells incubated with NIR QDs) (Fig 4D). It is important to mention that the scales for B16-Luc2 and HEK-Lux are different due to the intensity of the bioluminescence emitted by each cell types. The presence of NIR QDs in the specified wells was confirmed by the fluorescence images (Fig 4B).

**Fig. 4:**
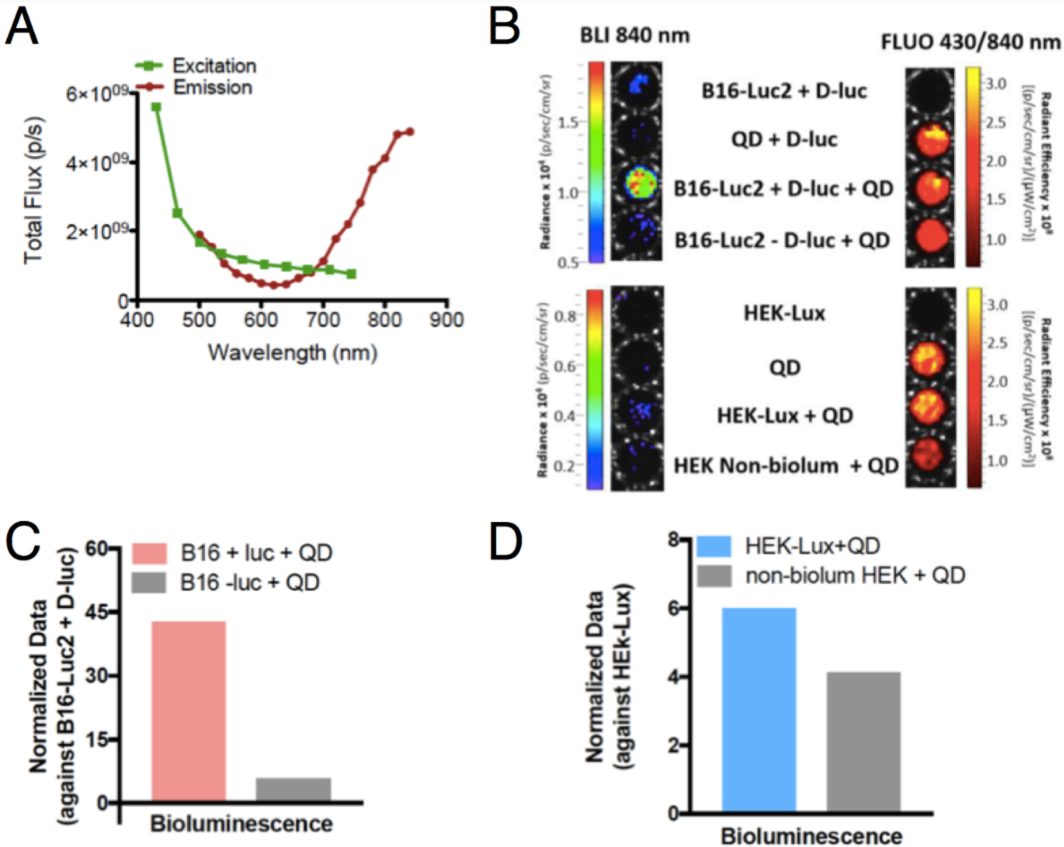
*In vitro* investigation of FUEL with NIR QDs. A) Excitation and emission fluorescence spectra of NIR QDs. Results are expressed as total flux (photons/sec) B) Bioluminescence (840 nm, exposure time of 60 sec (B16-Luc 2 cells) and 180 sec (HEK-Lux cells), as well as fluorescence images (excitation 430 nm, emission 840 nm and exposure time of 1 sec). C) Quantitation of bioluminescence signal emitted at 840 nm. Results are expressed as normalized SSMD values for B16-Luc2 cells (B16-Luc2 cells + D-luciferin used as control) or D) HEK-Lux (or non-bioluminescent HEK used as control). n=8 (except for HEK-Lux + QD − n=6).

We next investigated the ability of FUEL to red-shift tumour emission at the NIR QDs wavelength, enhancing the detection of tumour at red range wavelengths. Mice bearing B16-Luc2 tumours were imaged after the i.p. administration of D-luciferin to evaluate the background signal at 840 nm (−QD/+luciferin) (Fig 5A). After the intratumoral injection of NIR QDs (+QDs/+luciferin), we observed a drastic increase in the bioluminescence signal at 840 nm, confirming the presence of FUEL and its ability to enhance tumour detection at 840 nm by red shifting the light emission. Fluorescence imaging confirmed the presence of NIR QDs in the tumour sites and bioluminescence imaging in open filter shows that both right and left tumours were bioluminescent upon the administration of D-luciferin. No signal was observed in the absence of the substrate (-QD/-luciferin and +QD/-luciferin).

**Fig. 5:**
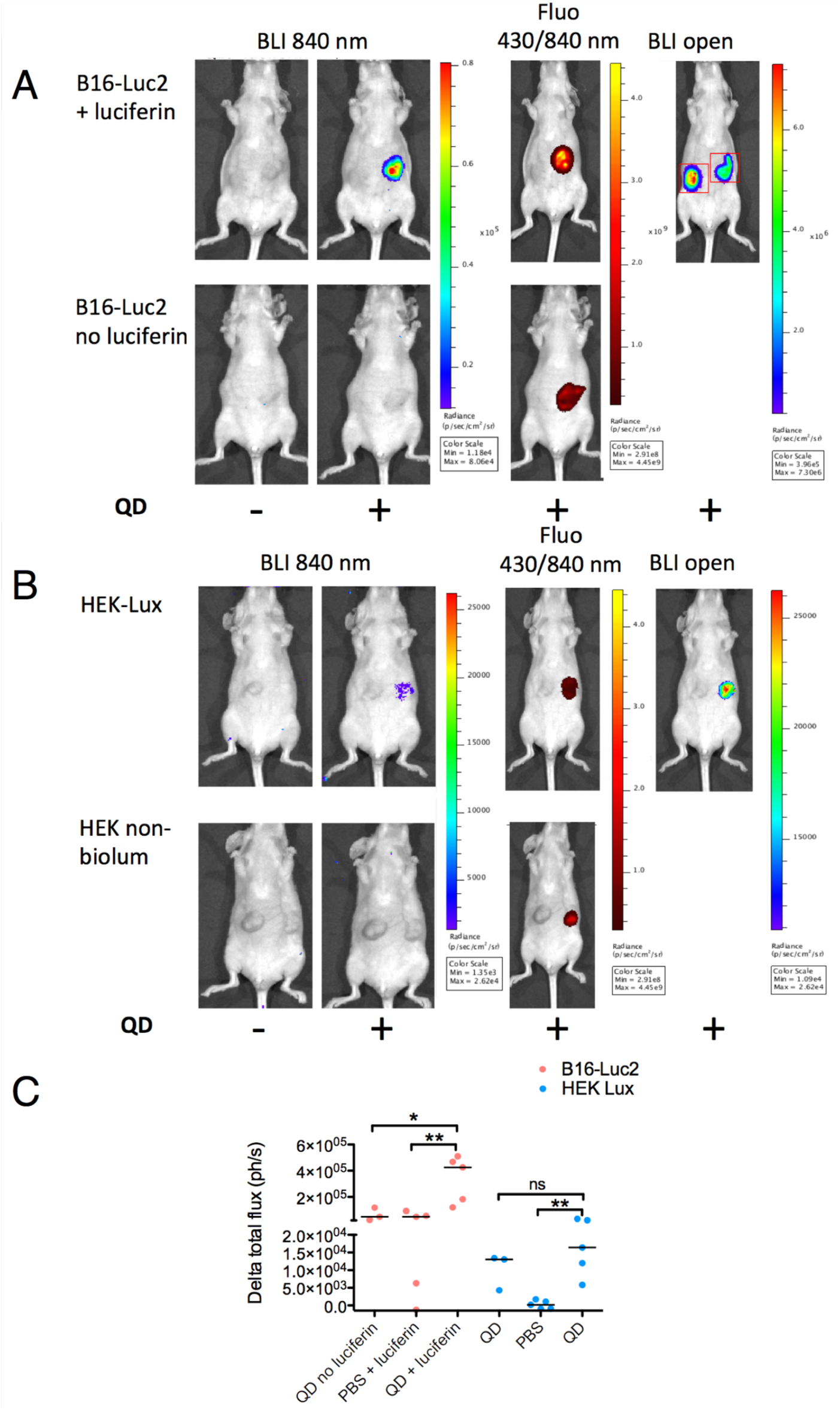
*In vivo* evaluation of FUEL. Bioluminescence imaging at 840 nm of B16-Luc2 (A) or HEK-Lux (B) tumours prior (left image) or after quantum dots injection in the right tumour (2^nd^ image left). Fluorescence images and bioluminescence in open mode are shown on the right. 840 nm bioluminescene images of control without luciferase for B16-Luc2 Cells (A) or non bioluminescent HEK cells (B) are shown in the second row. C) Quantitation of FUEL phenomenon. ROIs were determined as shown in the image of Figure 5A. Results express the delta between total flux (photons/sec) in the ROI of the left tumour of the mice post NIR QDs injection and prior to the NIR QDs injection, n=3 (negative control groups), n=6 (B16-Luc2) and n=4 (HEK-Lux). *p<0.05* was considered as significant: *\*p<0.05* and *\*\*p<0.001*.

FUEL efficiency was also investigated in HEK-Lux-induced tumour model. Bioluminescence signal at 840 nm post-intratumoral administration of NIR QDs was stronger than pre-injection (−QD/HEK-Lux vs +QD/HEK-Lux, Fig 5B and 5C). NIR QDs administered into non-bioluminescent HEK293 tumours showed bioluminescence signal statistically similar to HEK-Lux tumours with NIR QD.

In summary, we have shown that both tumour models undergo a red shifting in their emission via FUEL, where the red-shifting emission strongly depends on the optical emission properties of the tumours and the quantum yield of the near-infrared emitting fluorescent probe.

## Discussion

The development of new techniques for detecting tumours in an accurate and simple way is vital to support the search for new therapies in oncology. In this study, we used two different bioluminescent tumour models to demonstrate for the first time, that the FUEL process can be used *in vivo* to red-shift bioluminescence tumour emission and enhance the detection of tumours.

Herein, we established two murine models of tumours to investigate FUEL. One of the models was xenogeneic and made use of human (HEK-Lux) cells, an autobioluminescent cell type [5]. The second model was syngeneic, induced by B16-Luc2, a murine melanoma cell type expressing the enzyme luciferase frequently used in preclinical oncology [27]. While B16-Luc2 tumour growth and their bioluminescence signal showed the same profile, HEK-Lux cells initially presented a high bioluminescence activity immediately after the subcutaneous injection before showing a marked decrease of this activity the following day. We believe that these cells needed to adapt to the new environment before propagating and forming the solid tumour. After this latency period, the tumours reached the maximal volume that corresponded with the second peak of bioluminescence emission.

Each of the developed models has advantages and disadvantages with regard to FUEL applications. HEK-Lux cells have the enormous advantage of being autobioluminescent due to its constitutive expression of the bacterial lux operon thus enabling convenient image acquisition without having to consider the biodistribution kinetics of exogenously added substrate *in vitro* or *in vivo*[5] as for the B16-Luc2 cells [20]. This required substrate injection is a limitation since the time between substrate injection and imaging needs to be strictly controlled to achieve reproducibility in the data, mainly when acquiring images using different emission filters before and after the injection of NIR QDs. In addition, melanin production by the B16-Luc2 cells might be a concern for this type of imaging. However, we observed that melanin expression becomes significant only 2 weeks after subcutaneous injection, after we performed our experiments, and that these cells are indeed suited for FUEL imaging (Fig. 5).

FUEL is a phenomenon that allows the red shifting of the light, enhancing the detection of bioluminescent tumours because of the reduction of tissue absorption and scattering of blue/green light. One of the requirements for effective FUEL is that the fluorophore should have a large Stokes shift, determining the requirement of an ideal bioluminescent emitting source at approximately 500 nm [15, 16]. In this context, the wavelength of the maximal bioluminescence emission peak of HEK-Lux cells would be another advantage over B16-Luc2 cells regarding FUEL. Indeed, HEK-Lux cells emit luminescence at a maximum peak of 490 nm [5]. By contrast, B16-Luc2 cells have a maximum emission peak at 600 nm. In our case, we were still able to observe FUEL with B16-Luc2 because we used NIR QDs, which have a large absorption range. Furthermore, B16-Luc2 cells showed much stronger bioluminescence signal intensity in comparison to HEK-Lux cells, requiring shorter exposure times during imaging and overall higher FUEL efficiency. Our results show that even if HEK-Lux cells have a more appropriate maximum emission wavelength to excite NIR QDs than B16-Luc cells, due to their lower luminescence intensity, the red-shifting emission is not optimal. Indeed, if we focus on the maximum emission wavelength of both cell types, 500 nm and 600 nm for HEK-Lux cells and B16-Luc2 cells respectively, NIR QDs absorb 4 times less at 600 nm than at 500 nm (Fig 4A). However, the emission of B16-Luc2 cells is about 800 times higher at their maximum emission wavelength compared to HEK-Lux emission at their maximum emission wavelength (when the same number of cells are compared). Even at 500 nm, B16-Luc2 emission is 14 times higher than HEK-Lux cells, for the same number of cells (Fig 4B). These results highlight the fact that FUEL efficiency is controlled by a combination of both luminescence spectrum and intensity and acceptor absorbance properties. NIR QDs have many advantages for FUEL applications; namely high excitation coefficient and photoluminescence quantum yield. Moreover, this specific type of NIR QD has been shown to provide a lower *in vivo* toxicity compared to classical NIR QDs mainly because they are not composed of heavy metals [28]. In addition, FUEL efficiency also depends on the imaging conditions. The emission filters used in this study have a 20 nm bandwidth, which limits the imaging of the red-shifted emission photons. Using larger emission filter bandwidth or long pass emission filter should significantly improve the FUEL efficiency.

Our results suggest that FUEL can be used to enhance the detection of deeper and metastatic tumours by red-shifting their emission. As an optical method, FUEL has the significant advantage of requiring affordable imaging systems and facilities [3] that are extremely valuable in preclinical research. However, the experimental conditions of FUEL phenomenon for detecting tumours warrants some improvement and characterization to be fully suitable for enhanced detection of deeper tumours *in vivo*. Several factors mainly need to be taken into account: the biodistribution of the QDs within the xenograft, considering the tumour heterogeneity, and the fact that the tumour micro-environment could affect both luciferase enzymatic efficiency and fluorophore quantum yield, and consequently overall FUEL efficiency. The enhanced permeability retention (EPR) effect exhibiting by tumours as a result of leaky vasculature, could favour the retention of nanoparticles [29]. An effective EPR effect is strongly dependent on the size of the nanoparticles, their surface chemistry and the type of tumour. For instance, the accumulation and distribution of micelles of various sizes was not substantially affected by the size in a colon adenocarcinoma (C26) model, while size prove to be important in a human pancreatic adenocarcinoma (BxPC3) [30]. Positively charged nanoparticles have been shown to have shorter circulation half-life, but enhanced internalisation due to their adsorptive interaction with the cell membrane. Interestingly, Yuan *et al*. demonstrated the enhanced tumoral retention of zwitterionic nanoparticles with switchable charge based on environmental stimulus [29, 31]. In our study, the i.v. injection of NIR QDs 0.05 nmol in our models did not result in tumour retention, suggesting the absence of the EPR effect under our experimental conditions (Supplementary Figure 1). We have shown that both tumour models are similarly vascularised, which allows us to suggest that, with some improvement in our experimental conditions, NIR QDs could reach the tumours via i.v. administration. One alternative would be the targeting of tumours by coupling the nanoparticles with antibodies or peptides. RGD (Arg-Gly-Asp) is a triple-peptide motif that has affinity binding to the integrin α_v_β_3_, which is highly expressed in neovasculature and many tumour lines [32] and nanoparticles coupled to RGD have been shown to target tumours and improve their visualization [33]. Antibody-coupled nanoparticles have also been used for specific targeting of the tumour in preclinical imaging. NIR QDs or iron oxides nanoparticles coupled to anti-HER2 showed high specificity in targeting subcutaneous ovarian and prostate xenografts [34]. The NIR QDs used here could also be conjugated to antibodies and/or targeting peptides like RGD to ensure accumulation in tumours and provide more suitable experimental conditions to detect metastasis and deep tumours. In summary, we have shown the development of two different tumour models and FUEL ability to red shifting their emission. With further improvements, this optical method could offer an attractive alternative for detecting smaller and deeper tumours.

## Acknowledgments

We would like to thank Marie-Anne Nicola, Pascal Roux, Audrey Salles (Imagopole), Patricia Flamant (Platform of Histology), Mai Ban, Naoko Furusawa and Yasushi Nakano (Konica Minolta) who shared their insight and technical expertise, Biliana Teodorova and Pierre Bruhns for providing the B16-Luc cells and Dan Close (490 BioTech) for providing the HEK-Lux cells. This study was supported by NEDO-Konica Minolta, and has received funding from the French Government’s Investissement d’Avenir programs Laboratoire d’Excellence “Integrative Biology of Emerging Infectious Diseases” (grant ANR-10-LABX-62-IBEID) and Infrastructure d’avenir en Biologie Santé “France Life Imaging” (grant ANR-11-INBS-0006). ISA was partially funded by a Winston Churchill Memorial Trust UK Travel Fellowship.

## Supporting information legends

**S1 Fig. Absence of QD EPR effect. A)** Fluorescence quantification in tumours and abdominal control regions. B) representative image of fluorescence 24hours post QDs injection showing tumours and control ROI.

